# Unraveling the Genetic Architecture of Cryptic Genetic Variation

**DOI:** 10.1101/033621

**Authors:** Anupama Yadav, Kaustubh Dhole, Himanshu Sinha

## Abstract

Cryptic genetic variation (CGV), hidden under most conditions, is the repressed genetic potential that can facilitate adaptation and evolution. The conditional manifestation of CGV has been claimed to explain the background dependence of causal loci as well as missing heritability. However, despite being proposed over 60 years ago, the genetic architecture and regulation of CGV and its contribution towards regulation of complex traits remains unclear. Using linkage mapping of mean and variance effects, we have identified loci that regulate phenotypic manifestation of standing genetic variation in a previously published dataset of biparental *Saccharomyces cerevisiae* population grown in 34 diverse environments. Based on our results we propose the existence of a gradient of buffering states for a population determined by the environment. Most environments show a tight buffering with additive, independent causal loci with little epistasis. However, as this buffering is disrupted, the underlying highly interconnected environment-specific genetic interactome is revealed such that each causal locus is a part of this network. Interspersed within these networks are generalist capacitors that regulate CGV across multiple environments, with one allele behaving as a capacitor and the other as a potentiator. Our study demonstrates the connecting link between architecture of hidden and visible genetic variation and uncovers the genetic networks which potentially underlie all complex traits. Our study establishes CGV as a significant contributor to phenotypic variation, and provides evidence for a predictable pattern underlying gene-gene and gene-environment interactions that can explain background dependence and missing heritability in complex traits and diseases.

**SUMMARY:** The phenotypic effects of cryptic genetic variation (CGV) are mostly hidden and manifested only under certain rare conditions and have the potential to facilitate adaptation. However, little is understood about its genetic regulation. We performed variance QTL mapping to understand the regulation of phenotypic manifestation of standing genetic variation in a biparental yeast population. We propose a model describing the connecting link between visible variation and CGV. We identify generalist capacitors and environment-specific networks that potentially underlie all phenotypes. This fresh approach of mapping causal loci can solve the long-standing mystery of missing heritability in complex traits and diseases.

## INTRODUCTION

Cryptic genetic variation (CGV) is a heritable variation that is phenotypically inactive or hidden under most conditions, but manifests phenotypically only under certain, often rare, genetic and environmental perturbations (Gibson and Dworkin 2004). In simpler terms, it is bottled up genetic potential (Gibson and Reed 2008). While described in some studies as a separate and an inexplicable class of genetic variation, it essentially refers to gene-environment interactions, *i.e.,* genetic variants show their phenotypic effects only in certain environments or conditional neutrality, and gene-gene interactions, *i.e.* variants show their effects in the certain genetic backgrounds or epistasis. While the idea of CGV was proposed almost 60 years ago (Waddington 1956), for decades, both population genetic studies as well as evolutionary hypotheses have focused on the additive effects of mutations. Low resolution of mapping techniques curtailed the identification of large-scale gene-environment and genegene interactions. With recent advances in sequencing technologies and mapping resolution (Liti and Louis 2012; Siegal 2013), the prevalence of CGV in quantitative traits is being recognized. Environment specificity and epistasis identified in multiple studies to understand the architecture of complex traits in model systems, plant and animal breeding or to study environment and background dependence of human disease alleles, have established CGV as an extensive contributor to the genetic landscape of quantitative traits (Gibson 2009; Queitsch *et al*. 2012).

Introgression lines in classical systems like *Drosophila* and plants show that an allele can show a range of effects depending on the genetic backgrounds an allele is functional in (Gibson *et al*. 1999). Comparative studies in genetically accessible yeast have shown that the majority of loci that affect growth, an important fitness phenotype, have environment or genetic background dependent effects (Cubillos *et al*. 2011; Matsui *et al*. 2015). The importance of CGV in human traits is exemplified by increasing prevalence of diseases like type 2 diabetes. A highly heritable disease, type 2 diabetes, shows both population specificity and lifestyle dependence. Its high heritability is due to exposure of CGV by the modern culture (Gibson and Reed 2008; Gibson 2009).

While prevalence and impact of CGV is being demonstrated by increasing number of recent studies, its regulation is still unclear. Whether CGV is randomly distributed without any discernible patterns or is it controlled by specific genes or genetic networks is one of the key debates in the field, understanding of which will provide an ability to study the CGV landscape and predict its effects on diverse phenotypes (Hermisson and Wagner 2004). To address this question, it is important to understand how does CGV accumulate. Multiple evolutionary hypotheses and molecular explanations have been provided to explain the existence of CGV. Kimura’s neutral theory of evolution states that most mutations that accumulate in a population are phenotypically neutral (Kimura 1977; Kitano 2004). Waddington (1956) proposed an explanation for this neutrality by stating that if the mutations are disadvantageous to the optimum phenotype their effects will be hidden or dampened resulting in an invariant population phenotype, a process known as *canalization.* These dampening mechanisms could be altered upon genetic or environmental perturbations, thus revealing heritable variation which could facilitate the process of adaptation (Le Rouzic and Carlborg 2008; McGuigan and Sgrò 2009). The process of canalization, proposed by Waddington, invoked two types of genetic buffering mechanisms (Hartman *et al*. 2001; Burga *et al*. 2012) that contain the effects of CGV in stabilized, well adapted populations – generic buffering systems called *capacitors* (Rutherford and Lindquist 1998) and specific buffering systems consisting of *genetic networks* (Carlson and Doyle 2002). Capacitors are the genes that have the ability to suppress and hence store large amount of genetic variation. Upon specific environmental cues or genetic perturbations, these capacitors would, therefore, yield high CGV, that would affect all fitness phenotypes. In an adapting population, the released CGV can either be advantageous or detrimental (Orr and Betancourt 2001). However, in an adapted population, release of CGV would be detrimental since the population already exists at its phenotypic optimum (Barton and Turelli 1989; Hoffmann and Merilä 1999). As a result, such capacitors are proposed to have pleiotropic effects. Theoretically, such evolutionary capacitors should demonstrate storage of CGV as well as promote an increased ability to accumulate mutations or *mutational robustness* (Draghi *et al*. 2010; Fares 2015). The second type of genetic buffering could arise when stabilizing selection for a particular phenotype results in evolution of specific genetic networks, which buffer genetic variation in that particular phenotype (Masel and Siegal 2009; Paaby and Rockman 2014). These systems have different consequences on evolution as well as on molecular regulation of various traits.

Various studies have provided both direct and indirect molecular evidence for both these buffering mechanisms. Waddington’s initial studies in *Drosophila* lead to detailed investigation in the same system, which resulted in identification of genes with the ability to store CGV – *Ultrabithorax* (Gibson *et al*. 1999), EGFR (Dworkin *et al*. 2003) and, the most commonly studied gene, *HSP90* (Rutherford and Lindquist 1998). These genes result in revelation of cryptic variants with diverse effects on the phenotype upon their perturbation. Studies of Hsp90, across species, have demonstrated its generalized effect in regulating CGV in diverse phenotypes in yeast, worms, flies, plants and fish (Jarosz *et al*. 2010). Other more recently identified examples include *HZT1* (Richardson *et al*. 2013) and *IRA2* (Taylor and Ehrenreich 2015) in yeast. Alternately, molecular compensation through duplicate genes, redundant pathways (Wu and Lai 2015) and high genetic crosstalk have been proposed as specialized networks that contain CGV. A study showed that secondary enhancers of *Shavenbaby* are required to regulate variation in trichome patterning upon extreme stress in *Drosophila,* thus demonstrating a phenotype specific network (Frankel *et al*. 2011). Indirectly, high-resolution mapping of the cellular interactome (Boone *et al*. 2007; Li *et al*. 2010; Laufer *et al*. 2013) have revealed independent networks that are employed in specific traits and diseases, called modules (Vidal *et al*. 2011). This modularity may indicate phenotype specific genetic networks, which would have evolved in response to stabilizing selection to contain CGV in a specialized manner.

While CGV has been proposed to accumulate in a stabilized population and revealed in an adapting population, the accumulation of CGV and genetic buffering are not necessarily synonymous processes (Gibson and Reed 2008; Paaby and Rockman 2014; Siegal and Leu 2014). CGV can be a result of conditional effects of alleles that are maintained by mutation-selection balance and genetic drift across populations. These variants may have conditional effects on specific biochemical pathways, which differ in different populations (Chandler *et al*. 2013; Chari and Dworkin 2013; Chandler *et al*. 2014). Thus a large amount of such CGV can hence exist with no perturbations in the underlying robustness of the system, as conditional epistasis (Gjuvsland *et al*. 2007). This idea is supported by a lack of association between ability to release CGV and accumulate mutations *(mutational robustness).* While Hsp90 has been shown to facilitate release of hidden genetic variants (Jarosz *et al*. 2010; Rohner *et al*. 2013) that may eventually aid in adaptation (Orr and Betancourt 2001), it has not been demonstrated to support accumulation of novel mutations. The only study that directly tested the two phenomena, release of CGV and mutation robustness, compared the abilities of Htz1, the gene encoding histone H2A.Z (Richardson *et al*. 2013). While Htz1 showed high epistasis, CGV was released both in presence and absence of the gene, with no effect on accumulation of novel mutations. Additionally, even Hsp90 has been shown to act as both a capacitor (contain CGV) and as a potentiator (release CGV) (Cowen and Lindquist 2005). Genetic mapping both in functional and repressed Hsp90 states resulted in different, but equal number of genetic loci regulating yeast growth (Jarosz and Lindquist 2010). While such studies do not question the relevance of studying CGV to understand the genetic landscape of complex traits, they propose that loss of robustness is just one of the mechanisms that results in release of CGV (Gibson and Reed 2008; Paaby and Rockman 2014; Siegal and Leu 2014). However, the extent of overlap between the two phenomena is not clear. Additionally, the co-existence of generalist capacitors and specialized networks is not well understood, owing to exclusivity of the studies investigating these two phenomena. Are capacitors a part of specialized networks? Or is their existence independent of the networks active in different phenotypes? Several network based studies further challenge the evolution of such generalized capacitors, with an incentive to maintain phenotypic robustness, by showing that biological networks have evolved to maintain robustness such that multiple genes can behave as capacitors (Bergman and Siegal 2003). High epistasis and environment dependence of this epistasis further backs the case for CGV being a part of all the biological processes (Hermisson and Wagner 2004).

In this study, we have studied the phenotypic manifestation of standing genetic variation to address two key questions in the field. How common is regulation and release of CGV? Is CGV regulated by a few independent pleiotropic capacitors that function across environments or is it a property of certain environment-specific network modules? How do these generalized and specific forms of genetic buffering overlap?

We have used linkage mapping to identify loci that regulate CGV in a biparental yeast population grown in diverse environments. These genetically diverse parental strains (BY and RM11) have accumulated large number of polymorphisms over the course of their evolutionary trajectories (Liti *et al*. 2009; Bloom *et al*. 2013). Linkage mapping in such populations allows comparison of the effect of the two divergent alleles of a locus on thousands of genetic variants. While conventional quantitative trait locus (QTL) mapping identifies genetic loci, alleles of which have different effects on the population mean, their effects on population variance are often ignored (Rönnegärd and Valdar 2012). While differential mean of the two alleles would demonstrate the effect of the two alleles on the average phenotype, differential variance would convey that while the population is invariant or canalized in the presence of one allele, the other allele allows the phenotypic manifestation of diverse variants that results in a high variance. Therefore, differential variance represents differential ability to regulate CGV (Lempe *et al*. 2013). The term to describe these variance-regulating loci, variance QTL (vQTL) was first introduced by Rönnegärd and Valdar (2011). While most of these loci tend to be small effect when compared with conventional QTL (Shen *et al*. 2012), some large effect vQTL such as *MOT1* (Forsberg *et al*. 2015) and *nFT* (Lee *et al*. 2014) have been identified in plants. In addition, vQTL mapping has been proposed to be a predictor of prevalence of gene-gene interactions, further supporting differential regulation of CGV by these loci (Paré *et al*. 2010; Rönnegärd and Valdar 2012).

This differential variance between allelic effects is different from micro-canalization. Micro-canalization is another form of robustness that refers to phenotypic variation within two genetically identical individuals, or clonal variation (Bergman and Siegal 2003; Levy and Siegal 2008). Here, we have specifically investigated loci, which regulate suppression and revelation of heritable genetic variation.

We used variance QTL (vQTL) mapping to identify loci, which show the ability to regulate CGV across diverse environments. We further used covariance analysis to study the pleiotropy of their variance regulation and identified the revealed CGV through QTL-QTL and vQTL-vQTL interaction analyses. We identify a gradient of types of environments that determines the regulation of CGV within them. Most environments are tightly regulated such that the majority of the loci show additive effect without vQTL and epistasis interactions. Only in a few rare environments, this robust state is perturbed such that almost all loci regulate both mean and variance. Such environments reveal high number of epistatic interactions that regulate yeast colony size. It is in these environments, where we compared effect and abundance of generalist capacitors and specialized networks. We identify multiple capacitors that regulate CGV across multiple environments, one allele behaving as a capacitor and the other as a potentiator. These capacitors are separate from the core network, which is active independently in each of these extreme environments. We identified a single unique network for each environment. These networks maintain tight invariant phenotypes, which are perturbed only in certain allelic combinations, by using either of the following two approaches: maximalist phenotype or minimalist phenotype. These specialized networks probably employ these different capacitors to suppress CGV and as a result these capacitors release CGV, which either have positive effect or negative effect on the phenotype, depending on the kind of network. Our results argue that the regulation of CGV is a fundamental force in evolution of genetic networks underlying all complex traits.

## RESULTS

### variance QTL mapping identified loci that regulate CGV

QTL mapping allows comparison of effect of two alleles of a locus on a heterogeneous population. In an artificially generated biparental population, like the one used for this study, the majority of the alleles are present at an equal frequency. Therefore, while a difference in the mean of the two populations based on a marker demonstrates the effect of that allele independent of other loci, the difference in variance of the population carrying that allele is a representation of the effect of the allele on phenotypic manifestation of other polymorphisms, or CGV.

Using a previously published dataset (Bloom *et al*. 2013) of a recombinant haploid population generated from a biparental cross between a lab strain BY and a vineyard isolate RM11, we carried out linkage mapping to identify genetic loci, which showed an allelic difference on the mean (QTL) and variance (vQTL) of colony size variation. This mapping was done independently in 34 diverse environments, ranging from different carbon sources to oxidative and DNA damaging stresses. QTL were estimated using F-statistic comparisons whereas vQTL were using Brown-Forsythe (BF) test (Figure 1A, see Methods).

**Figure 1.**
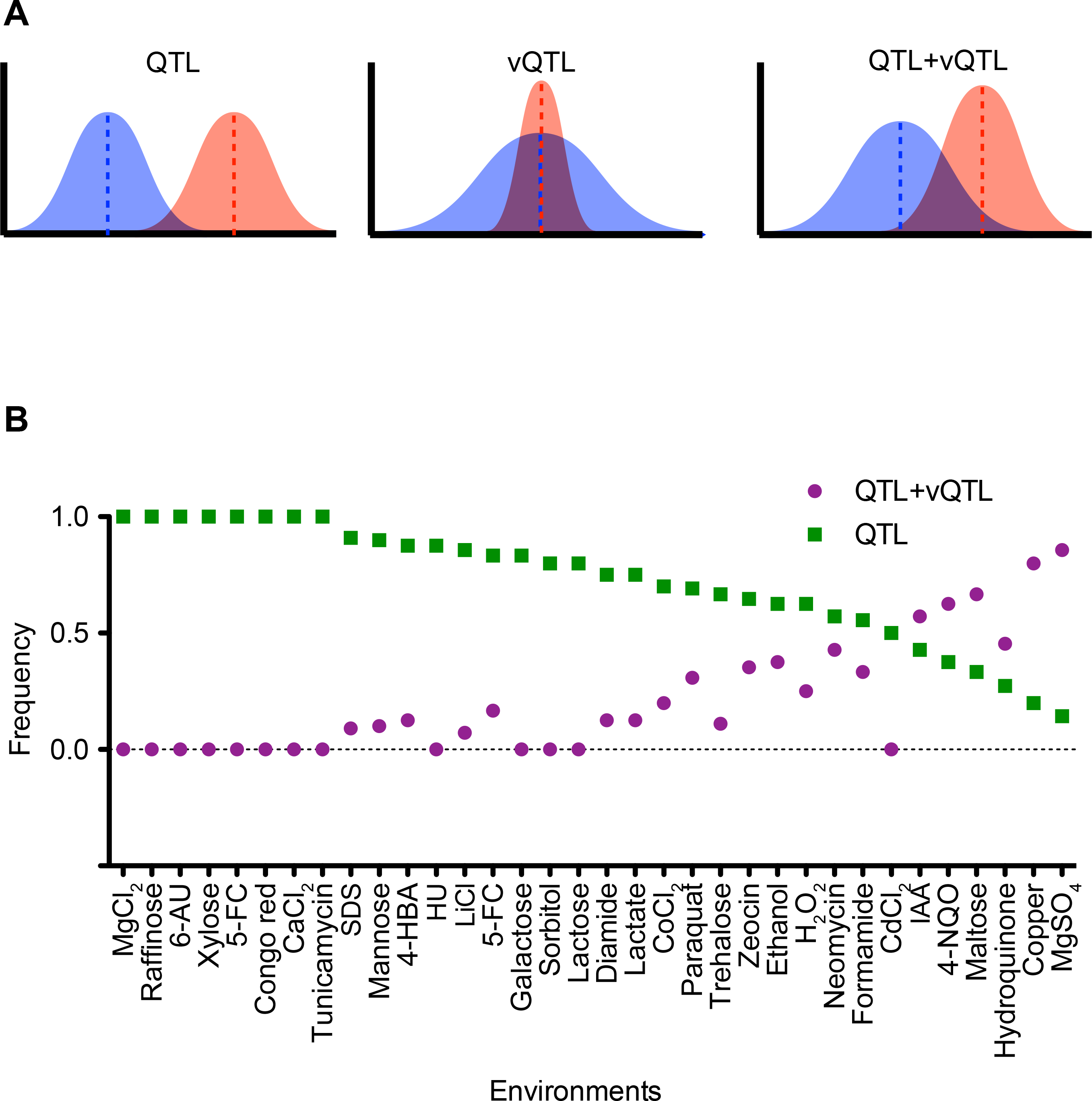
QTL and vQTL mapping. (A) Schematic showing three categories of QTL mapped. QTL has a significantly different allelic mean but a non-significant difference in allelic variance; vQTL has a non-significant mean difference but significantly different variance; QTL+vQTL has both significantly different allelic mean and variance. (B) Distribution of loci mapped in each environment as QTL (green) and QTL+vQTL (dark pink) in the segregating population. The y-axis is proportion of QTL or vQTL in each category. The x-axis is different environments arranged by decreasing proportion of QTL. Permutation value cut off < 0.01. See Table S1 for details.

By comparing the mean and the variance affects of the identified causal variants, we aimed to address the following main questions in this study. Is CGV pervasive in a population such that most loci show a basal difference in variance? Or do specific hubs, *i.e.,* capacitors exist that regulate the release of CGV across traits? And finally, what is the association between phenotypic robustness and CGV; if capacitors do exists, are they present in all environments such that their existence is decoupled from the state of phenotypic robustness of a trait. Or is the role of these capacitors revealed only in certain conditions in which robustness is perturbed?

Previous studies have mapped vQTL in segregating populations but their effect sizes were found to be small. Additionally, genetic studies claimed that many loci were capable of regulating phenotypic variance. To estimate the number of loci capable of affecting variance and their effect sizes, the loci were shortlisted separately based on significance and LOD score cutoffs (see Table S1). Based on significance of effects (P < 0.005), 44% loci behaved as only QTL and 16% as only vQTL across all environments (Table 1). A large proportion, 39%, of markers had a significant effect on both the mean and the variance. This indicated that more than half (55%) of the loci (including only vQTL and QTL+vQTL loci) affected variance. However, as previously observed, the differences in the population mean were more distinguishable than the differences in the population variance. Based on effect sizes (LOD score > 3, permutation *P* < 0.01), 70% of the loci behaved as only QTL, 6% as only vQTL and the remaining (24%) as both (Table 1). Thus while more than 50% of the loci had a significant effect on variance, the effect size was small in most cases. A high overlap was identified between loci regulating the mean and the variance, which indicated that release of CGV was associated with a shift in the population mean. Along with highlighting the ability of commonly studied QTL to regulate population means, this result emphasizes a specific directionality of release of CGV.

**Table 1.** QTL, vQTL and QTL+vQTL

| Category | No. of QTL | No. of vQTL | No. of QTL+vQTL |
| --- | --- | --- | --- |
| Based on $P < 0.005$ | 39 (16.6%) | 104 (44.3%) | 92 (31.1%) |
| Based on LOD $> 3.0$ | 19 (6.4%) | 212 (71.6%) | 65 (22.0%) |

If presence of capacitors and release of CGV is a common process, uncoupled from inherent robust or buffered state of the population in a given environment, then one would expect a uniform distribution of loci regulating variance across environments. Instead, we observed clustering among these loci regulating phenotypic variance along with mean (QTL+vQTL) in specific environments. A high negative correlation *(r^2^* = −0.9) was observed between ratios of loci classified as QTL and as QTL+vQTL across environments (Figure 1B). The figure shows that while the majority of environments show exclusive QTL effects, the enrichment of QTL+vQTL loci in certain environments demonstrates that most loci in these environments result in difference in variance along with the mean.

### Environment determines the release of CGV

A fundamental question associated with release of variance has been its effect on the average population phenotype, *i.e.,* whether release of variance will generate low or high fitness phenotypes, or both *(i.e.,* CGV will be released equally around the mean). A high number of QTL+vQTL and a few vQTL identified in our analysis established that the release of genetic variance by an allele was exclusively either advantageous or deleterious for the phenotype of the segregants (Figure 2A, see Table S1). While we cannot comment on the effect of colony size on fitness, we identified examples of loci that resulted in either only high or only low mean associated with release of variance indicating that loci were able to exhibit both capabilities (Figure 2A). In very few cases was the release of genetic variance by an allele spread equally around the phenotypic mean. One obvious question, from these observations, is which factors determine the direction of release of variance – does it depend on the locus or is it a property of the environment? Little association was observed between the mean and the variance in different pleiotropic loci across environments. For example, a chrXV (140,012) marker that had a significant effect on the mean and the variance in various environments showed no association between the two (Figure 2B, see Table S2), while the BY allele has a higher mean in MgCl_2_ and LiCl, and RM allele has a better mean in MgSO_4_; the RM allele has a higher variance in all these 3 environments suggesting poor correlation *(r^2^* = 0.3) between the mean and the variance for this particular locus (Figure 2B, see Table S2). However, a strong association between the mean and the variance of loci was observed within each environment (Figure 2B). In environment MgSO_4_, a strong positive correlation (*r*^2^ = 0.9) was observed such that an allele with a higher mean always had a higher variance independent of the mapped locus. Pair-wise correlation was calculated between the mean and the variance of all significant loci for each environment independently (see Methods). A poor correlation across the environments would have indicated that different loci resulted in random release of variance in both positive and negative directions within an environment. Interestingly, a significant correlation (*P* < 0.01) was observed in majority of the environments (26/33, Figure 2C). One half of the environments (13/26) showed a strong positive correlation (*r*^2^ > 0.5) indicating that release of variance was, on an average, advantageous for the phenotype. The other half environments showed a negative correlation (*r*^2^ < −0.5), *i.e.,* release of variance resulted in reduced colony size (Figure 2C). This meant that independent of the molecular nature of the regulatory locus or the hidden genetic variation, effect of release of variability on the population mean was strongly dependent on the environment.

**Figure 2.**
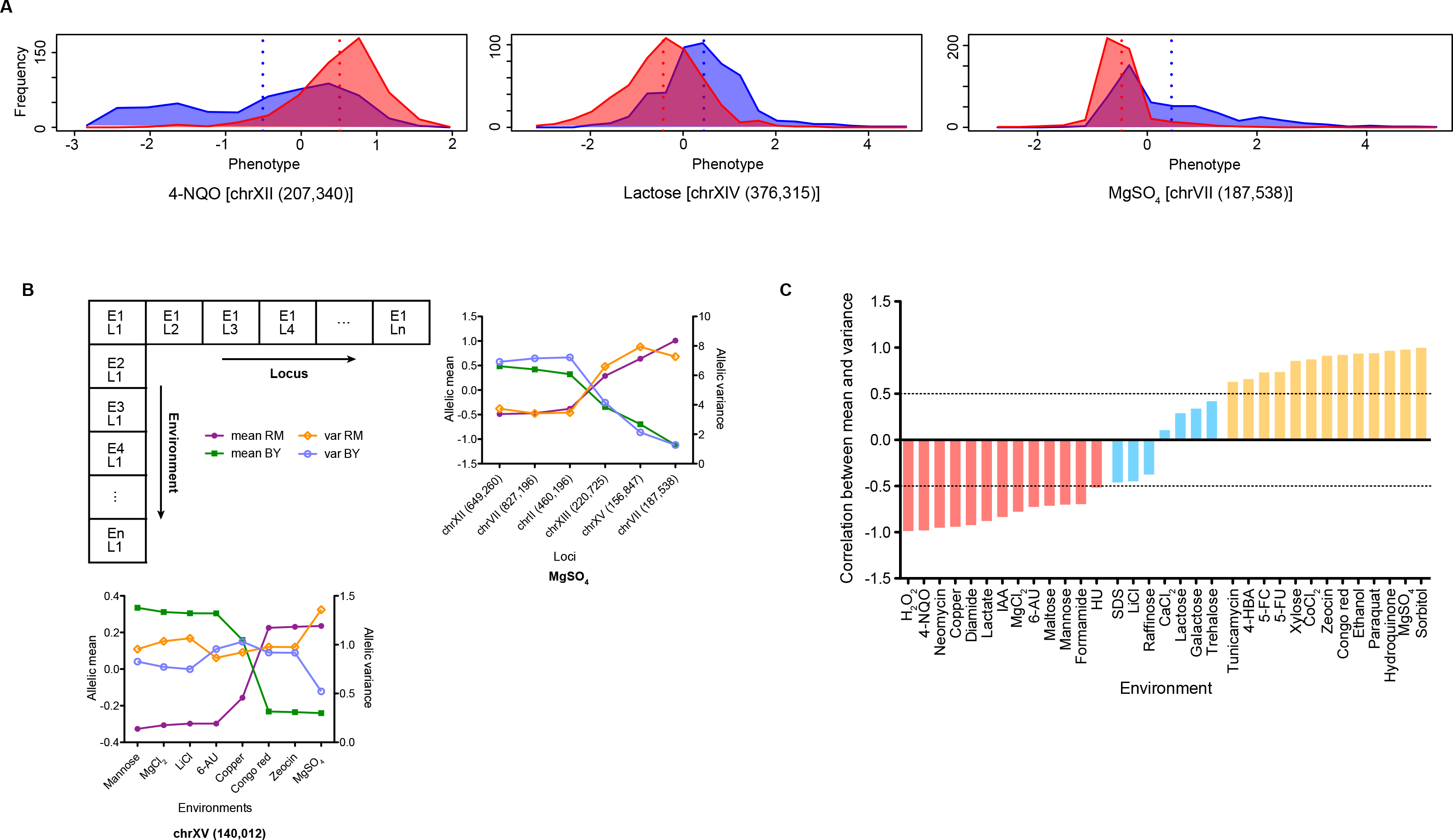
Environment and locus dependence of variance. (A) Representative frequency distributions of three loci showing directionality of release of variance. Blue distribution is of segregants with BY allele and red is for segregants with RM allele. 4-NQO [chrXII (207,340)] marker shows release of variance of the BY allele in the negative direction (mean RM > mean BY); Lactose [chrXIV (376,315)] marker shows equal variance of the two alleles; MgSO_4_ [chrVII (187,538)] marker shows release of variance of BY allele in the positive direction (mean RM < mean BY). The QTL are indicated as chromosome number followed by marker position in bp within brackets. (B) Environment and locus dependence of the mean and the variance of loci. The horizontal arm of the L-shaped grid represents multiple loci L1-Ln significant in environment E1. The corresponding graph shows the mean and the variance of BY and RM alleles of all markers (x-axis) in MgSO_4_. The vertical grid represents pleiotropic effect of the locus L1 in multiple environments E1-En and corresponding graph shows mean and variance of the marker chrXV (140,012) in various environments (x-axis). (C) Correlation between the mean and the variance of all QTL in each environment. Correlations less than −0.5 (red, *P* < 0.05) represent environments that have a negative association between the mean and the variance. Correlations more than 0.5 (yellow, *P* < 0.05) represent environments that have a positive association between the mean and the variance. Environments with no directional release of variance (correlations between ±0.5) are represented as blue. The y-axis is correlation between mean and variance; dashed lines mark ±0.5 correlation value.

This prepotency of the environment in determining direction of release of variance raised the possibility of environmental constraint on the nature of released CGV. Do same loci regulate the mean and the variance across different environmental categories (positively versus negatively correlated environments)? If yes, then do these alleles show similar or opposite effects on the regulation of variance between different categories? A comparison of QTL+vQTL effects of a locus across environments was limiting due to higher number of QTL effects than QTL+vQTL or vQTL effects (see Table 1 and Figure 2B). More importantly, alleles of a locus may show similar variance in an environment but behave differently across environments. Trait covariance measures how a population behaves across two environments (Haber and Dworkin 2015). A high covariance means similar phenotype, and similar identity of CGV released in this case, whereas a low covariance indicates differential phenotype of the segregants across the two environments. Therefore, while the two alleles can have similar variance, they may show differential release of CGV, across two environments (Figure 3A). BY allele of the chrXV (160,440) marker showed similar variance in Lactate (*σ*^2^ *=* 0.88) and Lactose (*σ*^2^ = 0.69), much like BY allele of marker chrXIV (681,897) in trehalose (*σ*^2^ = 0.77) and LiCl (*σ*^2^ = 0.86). However, the latter allele showed a much lower covariance (*Cov* = 0.13) than the former (*Cov* = 0.84). This indicated that while the RM allele allowed similar phenotypic manifestation of genetic variants in the population in both Lactate and Lactose, BY allele showed differential regulation of these variants across Trehalose and LiCl. Mapping covariance of loci across pairs of environments therefore, will identify loci exhibiting differential CGV across environments, and hence uncover potential capacitors. In addition, covariance mapping will also determine whether allelic bias exists between the ability to release variance, *i.e.,* does one allele always result in release of variance independent of the environment or this ability of an allele is determined by the environment in which the locus is present in. In other words, does an allele behave exclusively as a capacitor or as a potentiator, or can it behave like both depending on the environment? As an alternative to performing a genome-wide covariance analysis, we chose only those loci that had a significant effect (QTL, vQTL or QTL+vQTL) in more than one environment. Forty seven such loci were selected and their covariance was computed across these environmental pairs (see Methods, Table S3). Eighteen loci showed a significant difference in the covariance with 9 being significant across multiple (more than 15) environmental pairs (Figure 3B, Table S3). These 9 loci regulated covariance across multiple environments and therefore were identified as hubs of variance regulation, *i.e*., capacitors or covariance hubs. The same locus regulated covariance in environments from different categories where the mean and the variance were positively correlated and negatively correlated (Figures 2C, 3C, Table S3). Furthermore, within a locus, while either of the alleles had the ability to result in release of variance across environments, a significant allelic bias in this ability was observed. Among the 5 loci that regulated covariance in more than 15 environmental pairs, 4 had a significant allelic bias (Fisher Exact test *P* < 0.05, Figure 3B). This meant that while both the alleles of these loci are capable of functioning as capacitors, one allele tends to contain variance and the other results in its release in majority of the environments. In Figure 3C, Paraquat shows a positive correlation between the mean and the variance whereas Cu shows a positive correlation. However, BY allele of the chrXIII (45,801) marker shows reduced variance in both environments whereas the RM allele shows a higher variance. Hence alleles of a locus tend to behave exclusively either as capacitors or as potentiators, independent of the environment.

**Figure 3.**
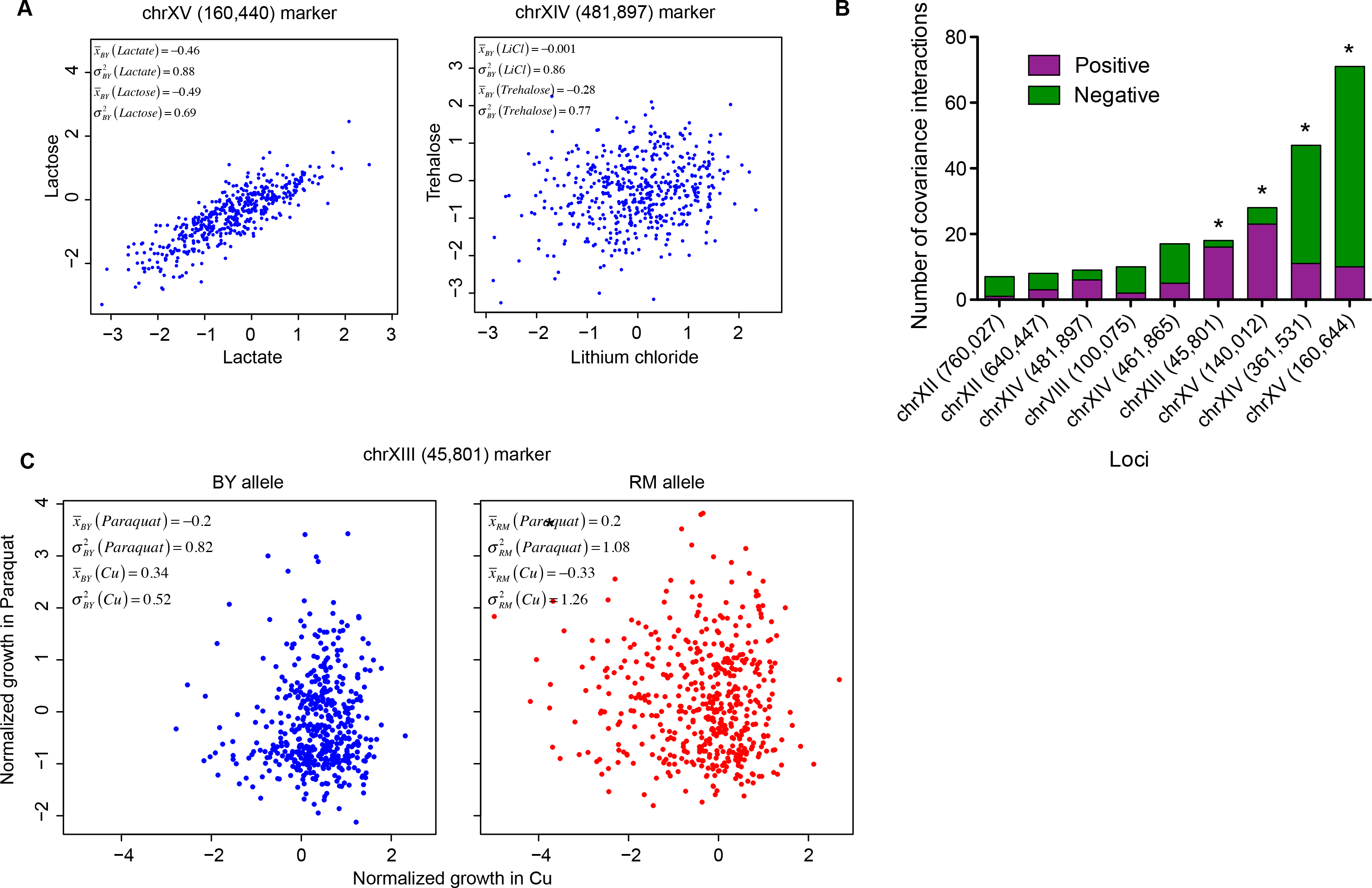
Allelic bias in across environment covariance. (A) Covariance of normalized growth phenotype of the BY (blue) allele of chrXV (160,440) marker in Lactose and Lactate. Covariance of normalized growth phenotype of the BY (blue) allele of chrXIV (481,897) marker in Trehalose and Lithium chloride (LiCl). The mean and the variance of each allele in each environment are indicated in the box. The QTL is indicated as a chromosome number followed by the marker position in bp within brackets. (B) Frequency distribution of number of positive and negative values of Deming regression t-test (see Methods) across pleiotropic covariance loci (x-axis). The t-test value is positive where Cov(BY) > Cov(RM) and negative for Cov(BY) < Cov(RM). The y-axis indicates number of environmental pairs across which the locus showed a significant difference in covariance. A significant Fisher Exact test (*P* < 0.05) is indicated by a star. The QTL is indicated as a chromosome number followed by the marker position in bp within brackets. (C) Covariance of normalized growth phenotype of the BY (blue) and the RM (red) allele of chrXIII (45,801) marker in Paraquat and Cu (Copper). The mean and the variance of each allele in each environment are indicated in the box. Paraquat shows a positive correlation between the mean and the variance whereas Cu shows a negative correlation (Figure 2C). The QTL is indicated as a chromosome number followed by the marker position in bp within brackets.

### CGV as a major contributor to gene-environment interactions, especially antagonistic pleiotropy

Environmental influence on the direction of release of variance (a better or a poorer phenotype) and high consistency of the regulation of variance by covariance-regulating pleiotropic hubs have a pronounced ramification on the role of CGV in regulating gene-environment interactions. While CGV explains the high conditional neutrality of genetic effects across environments, we demonstrate that the direction of release of CGV, as determined by the environment, has a strong effect on gene-environment interactions, especially antagonistic pleiotropy. Assume a following scenario where a locus regulates the mean and the variance across two different environments, *i.e.,* there is opposite correlation between the mean and the variance. Since there is a high consistency in regulation of variance, one allele will always result in high variance, while the other allele will always result in lower variance, and hence the high variance allele of this marker will have opposite effects on the mean thus resulting in antagonistic pleiotropy. For example, the locus chrXV (140,012), in Figure 2B, resulted in differential variance and mean in environments MgCl_2_, LiCl and MgSO_4_. Because LiCl and MgSO_4_ have opposite correlations for the mean and the variance, the consistency of RM allele in showing higher variance results in a trade-off in the overall mean effects. This phenomenon is clearly demonstrated by the example in Figure 3C, where BY allele of chrXIII (45,801) marker had better mean in Cu and poorer mean in Paraquat than the RM allele and hence showed antagonistic effects. Therefore, in this example, this antagonism was a result of the consistency of the RM allele in reducing variance and the BY allele in releasing variance. Thus, opposite correlation between the mean and covariance in Paraquat and Cu resulted in antagonistic effect of this locus. Therefore, CGV and its regulation contribute to gene-environment interactions in multiple complex ways.

### Two-locus mapping identifies the hidden variants

Identification of loci acting as hubs regulating covariance indicates their ability to function as capacitors. However, to conclusively determine their ability to regulate CGV, identification of the hidden variants is crucial. As previously stated, release of CGV is dependent on genegene interactions or epistasis between the stimuli (capacitor or genetic background) and the hidden genetic variants. The conventional use of QTL-QTL mapping is to identify epistatic interactions that contribute to variation in a population. We utilized QTL-QTL mapping to identify the genetic basis and patterns of regulation of released CGV. In addition, we adapted the QTL-QTL mapping technique to perform vQTL-vQTL mapping, which would allow us to identify the loci that show interactive effects of the allelic combinations on the variance of the population instead of the mean. We used QTL-QTL and vQTL-vQTL mapping to understand following points regarding the genetic basis and patterns of regulation of released CGV (see Table S4). Are most genetic interactions environment-specific, such that release of CGV is merely conditionally neutral and environment-specific epistasis? Or do discrete loci exist that behave as capacitors and release CGV across multiple environments? If the covariance hubs, identified previously, indeed function as capacitors, then they should show a high number of genetic interactions across environments. Are there genetic networks that maintain CGV in a population-specific and environment-specific manner? A perturbation of these networks, either due to a rare environment or genetic intermixing, would result in release of CGV. Such a scenario would highlight the role of such network(s) beyond regulation of CGV, in maintaining phenotypic robustness.

QTL-QTL and vQTL-vQTL mapping was carried out in an environment dependent manner within all loci that were significant as QTL, vQTL or QTL+vQTL in those environments. Total of 73 significant interactions (P < 0.1) were identified of which 18 (24%) were QTL-QTL, 33 (45%) were vQTL-vQTL and 22 (30%) were both QTL-QTL and vQTL-vQTL interactions (Table 2). This substantially high number of vQTL-vQTL interactions, in comparison to proportion of single vQTL (Table 1), indicated their role in contributing to population variation without demonstrating epistasis of the mean effects. Interestingly, amongst the loci showing QTL-QTL interactions, 10% had only QTL effects, whereas 80% were either vQTL or QTL+vQTL (Table 2). In addition, while there was a negative correlation between the total number of QTL in an environment and the number of QTL-QTL (*r*^2^ = −0.46) and vQTL-vQTL (*r*^2^ = −0.37) interactions, a strongly positive correlation was observed between number of QTL+vQTL and number of QTL-QTL (*r*^2^ = 0.64) and vQTL-vQTL (*r*^2^ = 0.65) interactions, indicating that the release of variance was a result of revelation of epistatic effects. In other words, a high number of QTL+vQTL effects is a robust predictor of epistasis in the trait.

**Table 2.** Two-locus interaction mapping

| Interactions ($P < 0.1$) | QTL-QTL | vQTL-vQTL | QTL-QTL+vQTL-vQTL |
| --- | --- | --- | --- |
|  | 18 (24.7%) | 33 (45.2%) | 22 (30.1%) |

### Identifying generalist capacitors and specialized genetic networks regulating CGV

By comparing the results of QTL-QTL and vQTL-vQTL interactions across environments, we attempted to distinguish between the three scenarios: environment-specific epistasis, capacitors and networks maintaining robustness. The loci showing interactions, within and across environments, was compiled. A locus that had interactions in two or less than two environments and single interaction per environment was considered as an environment-specific interactor; a locus with interactions in more than two environments was considered a capacitor; and a locus with more than two interactions within an environment was considered a part of a network active within that environment. A total of 73 interactions (QTL-QTL, vQTL-vQTL, QTL-QTL+vQTL-vQTL) were identified, which meant 146 interactors (see Table S5). Amongst these, 29 interactors were involved in environment-specific interactions, 55 interactors which attributed to 9 loci showed interactions across diverse environments, *i.e.,* behaved as capacitors. The remaining 62 showed multiple environment-specific interactions. Of the 55 markers that behave as capacitors, 62% marker showed interactions that had a difference in variance (vQTL-vQTL or vQTL-vQTL+QTL-QTL). Interestingly, of the 62 markers, which showed environment-specific interaction networks, 85% showed a difference in variance, indicating that these capacitors and networks regulate CGV possibly through higher order interactions. Only 2 loci were involved in multiple interactions across environments (chrVIII and chrXIII). These results showed that environment-specific interactions were rare, and loci which form environment specific networks are different from the ones which behave as capacitors.

Epistasis across traits follows a pattern where interaction hubs regulate CGV across multiple environments. Amongst the 9 interaction hubs identified, 7 also behave as covariance hubs indicating that loci that regulate variance across multiple environments do so by interacting with different loci in different environments (Figures 4A, 5A, Table S3). When combined with consistency in direction of covariance, these results demonstrate how CGV is released majorly in the presence of one allele of the capacitor and not the other (Figure 4A). Different genetic variants are released across different environments. The Figure 4A shows that BY allele of the chrXIV (368,185) marker has a higher variance in both 4-HBA and Galactose and the RM allele has a lower variance. In 4-HBA, the chrXIII locus shows a larger effect in the presence of the BY allele than the RM allele (Figure 4B). Similarly, in Galactose, chrXV locus shows a stronger effect on the phenotype in the presence of BY allele than the RM allele (Figure 4C). These results demonstrate how the covariance hubs result in release of differential CGV across different environments.

**Figure 4.**
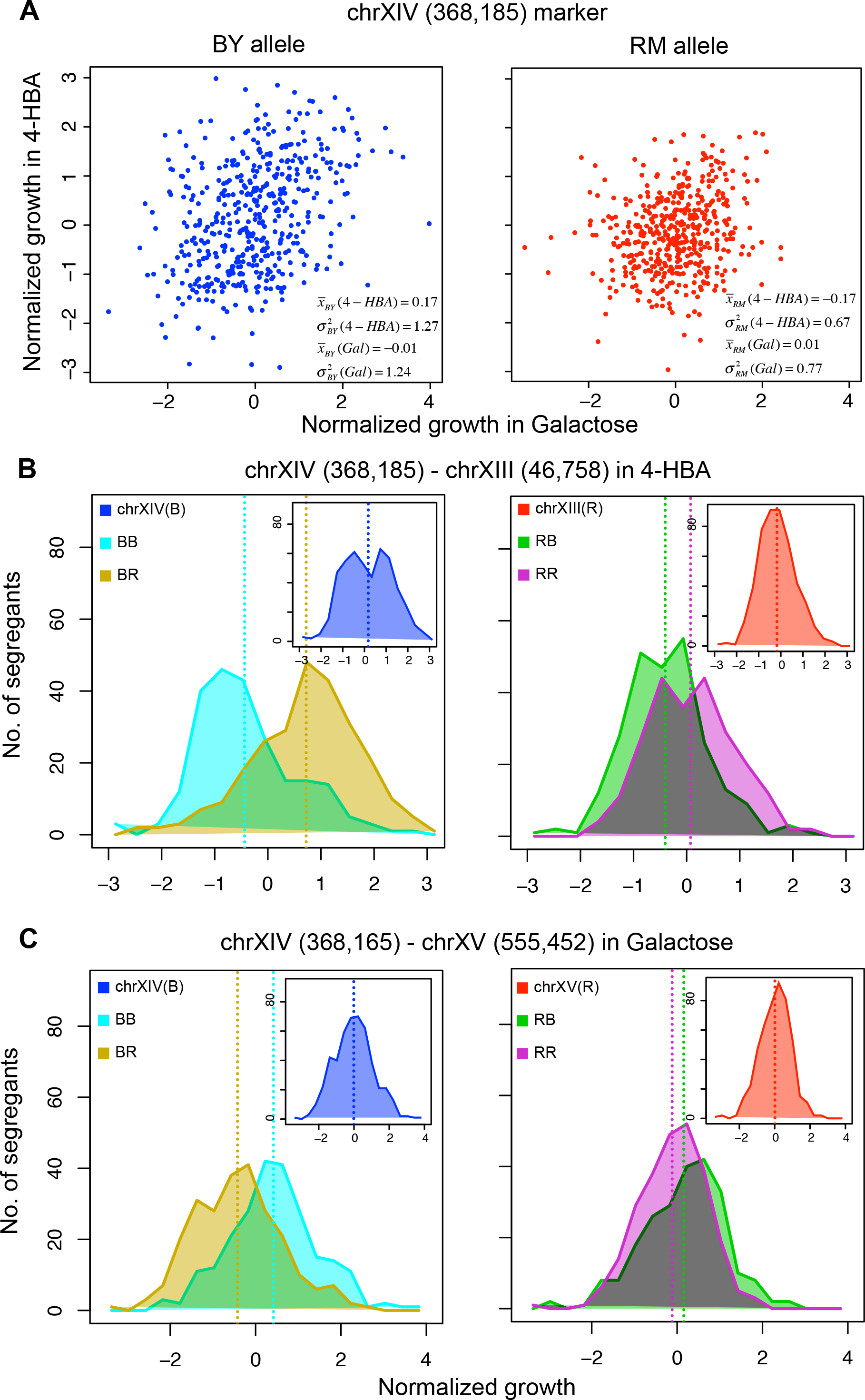
Release of CGV through QTL-QTL interactions. (A) Covariance of normalized growth phenotype of the BY (blue) and the RM (red) segregants for chrXIV (368,185) marker in 4-HBA and Galactose. The mean and the variance of each allele in each environment is indicated in the box. The axes are normalized growth of segregants in the two environments indicated. (B) QTL-QTL interaction between chrXIV (368,185) - chrXIII (46,758) in 4-HBA. (C) QTL-QTL interaction between chrXIV (368,165) - chrXV (555,452) in Galactose. The QTL is indicated as a chromosome number followed by the marker position in bp within brackets. For (B, C), the x-axis is normalized growth of segregants in the environment and the y-axis of number of segregants. Dash lines in segregant distributions (B, C) indicate the means of the distributions. The biallelic marker segregant distributions (in the QTL marker order written above the plots) are indicated as BB (light blue), BR (light brown), RB (dark green) and RR (purple). Inset plots show the average distributions of the first marker (BY (blue) and RM (red) alleles). See Table S4 for details.

Sixty six markers showed environment-specific networks in 12 environments (see Table S5). Some of these networks were denser than others. These networks showed high interconnectivity and the loci in these networks showed QTL+vQTL effects. Additionally, the interactions showed both QTL-QTL and vQTL-vQTL effects, *e.g.* a network of 4 loci present in Indoleacetic acid (IAA) where all loci show significant interactions with each other (Figure 5B, C). As demonstrated by the dot plots, the allelic combinations differed in mean as well as variance (Figure 5B). As shown above, the differential variance between alleles of a locus is deterministic of allele-specific revelation of effects. This finding (derived from many such examples as one shown in Figure 5, also see Figure 6B) can be extrapolated to state that loci showing vQTL-vQTL interactions harbour higher degree interactions within them, which results in differential variance of the allelic combinations. Indeed, as shown in the Figure 5B, vQTL-vQTL interactions occured when 3 out of 4 biallelic combinations exhibited lower variance, higher mean phenotype, whereas one combination resulted in increase in the variance and reduction in the mean. Similar networks were present in other environments as well, with these environments demonstrating a range of connectivity patterns. While all 4 loci showed similar effects in IAA, networks in other environments have loci that act as hubs showing higher interactions than other members of the network (see Figure 6B). A consistent property of an environment containing such networks is that all loci showing an effect on the phenotype in these environments are a part of this network. They may be present as highly interacting hubs or as interactors of one of these hub loci (see Figure 6B). Additionally, most of the loci of these networks are exclusive to each environment (see Figure 6B). This indicates a revelation of underlying genetic networks in these environments, with rare additive effects of loci. A tempting possibility is that this dense genetic interactome underlies all traits, but is revealed under specific genetic and environmental perturbations.

**Figure 5.**
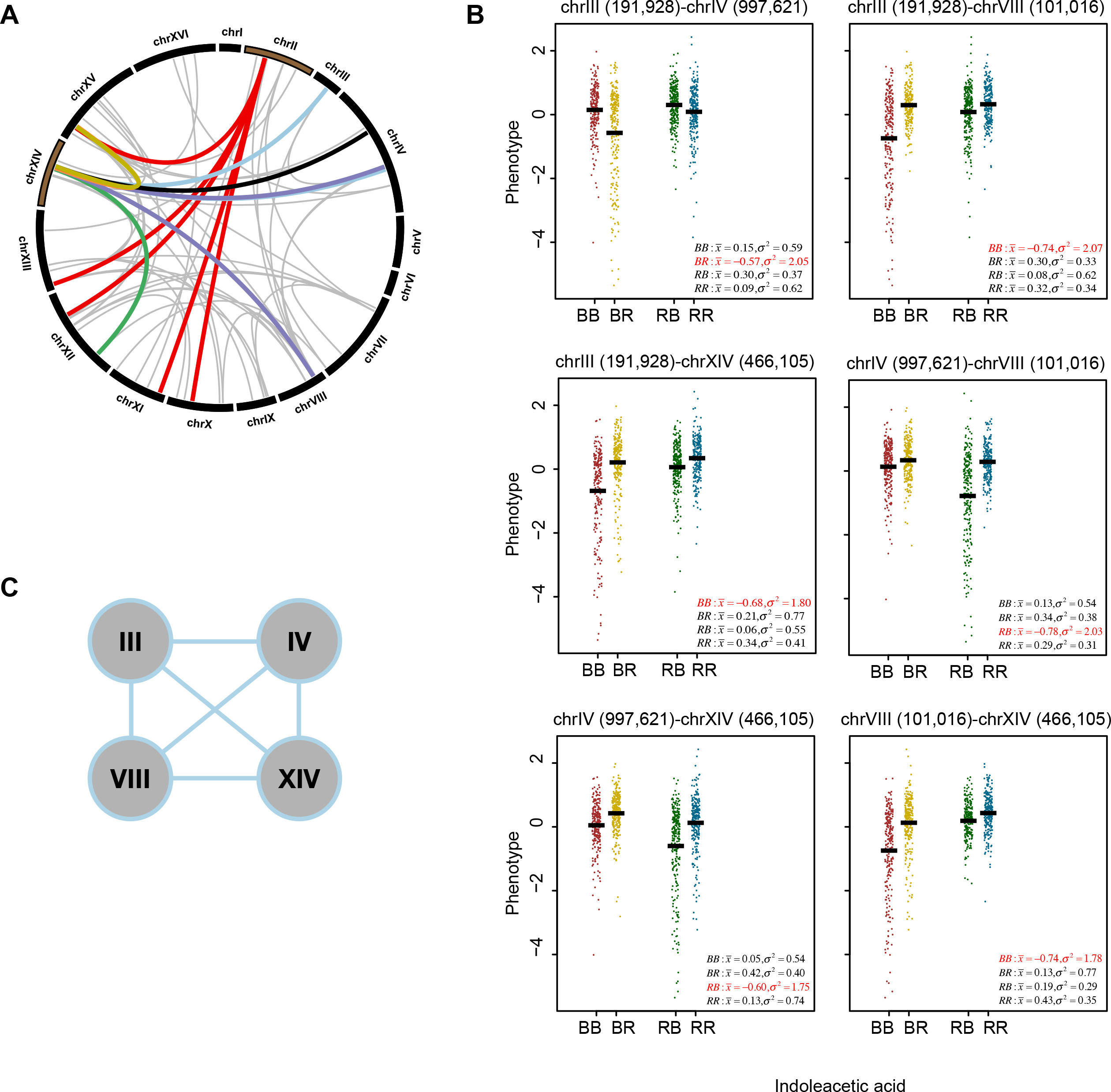
QTL-QTL interaction hubs and networks. (A) QTL-QTL interactions between various markers shown as connected links. The chrII (245,879) marker (red) has multiple QTL-QTL interactions (deep red) for growth in same environment (Congo red). The chrXIV (466,590) marker (green) has 6 environment-specific QTL-QTL interactions (4-NQO = dark green, Formamide = deep purple, Indoleacetic acid (IAA) = light blue, Lithium chloride (LiCl) = deep orange, Trehalose = deep yellow, Xylose = orange). Other QTL-QTL interactions are indicated as light grey links. Genomic interaction maps made using Circos (Krzywinski *et al*. 2009). See Table S4 for data. (B) Scatter plots showing examples of QTL-QTL interactions of four markers [chrIII (191,928), chrIV (997,621), chrVIII (101,016), chrXIV (466,105)] in Indoleacetic acid (IAA). The biallelic marker segregant distributions (in the QTL marker order written above the plots) are indicated as BB (red), BR (yellow), RB (green) and RR (blue) on x-axis. The mean and the variance of each allelic pair is indicated in the box with allelic pair with most variance indicated in red. The y-axis is normalized growth phenotype. (C) Schematic representation of genetic network of four loci, indicated in (B) above, showing QTL-QTL interactions in Indoleacetic acid.

## DISCUSSION

The phenomenon of CGV was first proposed over 60 years ago (Waddington 1956). While its existence has been demonstrated in various systems across taxa, the architecture of cryptic genetic variation and its effect on the genetic regulation of complex traits are still unclear (Siegal and Leu 2014). Several questions surround the understanding of regulation of CGV.

How is it regulated – by generalist pleiotropic capacitors or by phenotype-specific networks? What effect does it has on the phenotype, in other words, what is the distribution of deleterious versus advantageous CGV? Can the effect of release of CGV on the phenotype be predicted? Is the architecture of the hidden genetic variants different from that of the additive loci (Gibson and Reed 2008)? Studies have demonstrated both detrimental and beneficial effects of CGV on various phenotypes (Jarosz *et al*. 2010). In addition, contribution of various phenomena *viz.* robustness, canalization, buffering and capacitance to CGV, further complicates its mechanistic understanding (Gibson and Dworkin 2004). We used an integrative approach of mapping loci regulating mean, variance and covariance and their epistatic regulation to understand the underlying genetic framework of CGV. Comparing genetic regulation of multiple environments allowed us to comprehensively study the regulation of CGV and the integrated co-existence of conventionally identified loci and CGV. We propose the following model based on our results (Figure 6A).

**Figure 6.**
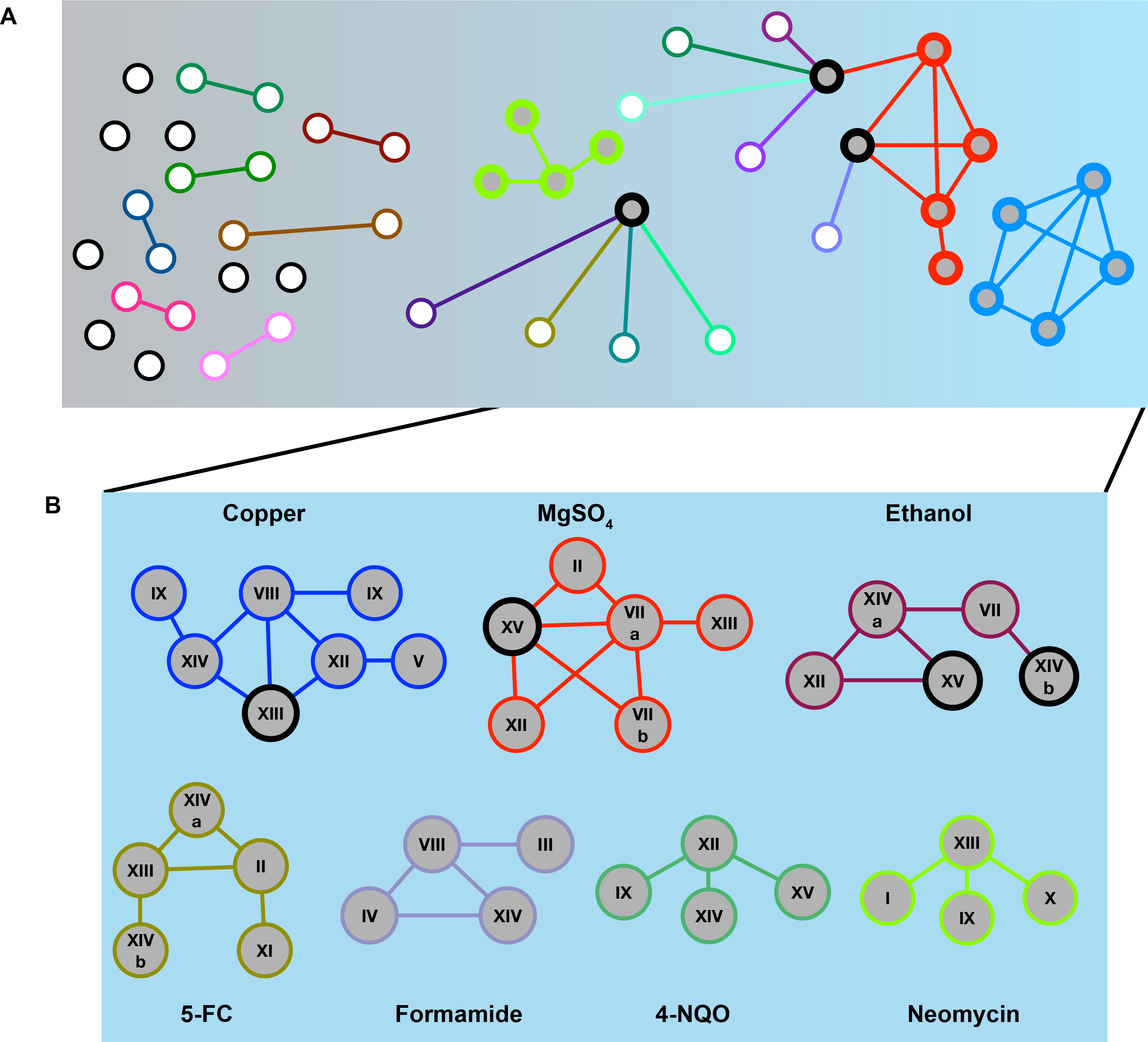
Genetic architecture of CGV. (A) A model showing gradient of phenotypic buffering from gray (left side) to blue (right side). Different environments fall in different positions along this gradient. The left side indicates robust buffering in the population such that most loci have additive independent effects with a few epistatic interactions. On the right side, are populations with revealed CGV that exposes the underlying genetic networks present in different environments. Different environments are represented by different colors. The white nodes refer to independent additive loci, gray nodes refer to loci which are a part of different genetic networks. The gray nodes with thick borders denote capacitors. Most of the environments fall on the left side of the gradient such that the majority of the QTL identified in different studies are additive with a few epistatic interactions. As the underlying buffering state of the population gets perturbed, due to either genetic or environmental perturbations, it moves towards the right side of the gradient. This perturbation reveals the underlying genetic network, which regulates phenotypic variation such that all loci are part of this highly interconnected network. These networks are environment-specific interspersed with generalist capacitors which regulate buffering and epistasis across different environments. (B) Examples of specific genetic networks identified in different environments using QTL-QTL and vQTL-vQTL interaction mapping. The potential capacitors identified through covariance mapping (see Figure 3B) are highlighted as nodes with black outlines. See Tables S4, S5 for details.

A population can exist in different states based on its genetic diversity and the surrounding environment. While we do not understand the underlying mechanistic differences between these states, they appear to be a gradient of the internal buffering states or phenotypic robustness (of heritable variation) of the population. Towards the left end (gray) are states where genetic variation is majorly additive in nature, with a few epistatic interactions such that most loci have independent effects with difference in the population mean. Some loci do show pleiotropic effects in these states but only affect the mean in different environments. For the biparental population used in this study, majority (more than half) of the environments are confined to this side of the spectrum. This result is substantiated by mapping of loci with predominantly additive effects and limited identification of epistasis in different studies (Carlborg and Haley 2004; Bloom *et al*. 2015). Additionally, this explains the low effect of variance QTL identified in multiple studies. Moving towards the right side (blue)along this gradient, the population state gets altered such a way that the underlying genetic interconnectivity, and hence CGV, gets revealed (Carlborg and Haley 2004; Mackay 2014; Siegal and Leu 2014; Fares 2015). This perturbation appears to happen in rare environments (Kitano 2004) and uncovers both kinds of genetic buffering mechanisms – capacitors, which regulate CGV across multiple environments and environment-specific genetic networks. Results from a recent study, which has identified capacitors and potentiators of bristle numbers in *D. melanogaster*, support our model, thus suggesting its universality (Takahashi 2015). Not just regulation of CGV, but these two mechanisms together form the core genetic regulation in these environments (Félix and Barkoulas 2015). The unusualness of these environments provides support for the Highly Optimized Tolerance (HOT) hypothesis, which states that systems evolved to be robust against most common perturbations, show high vulnerability to rare conditions (Carlson and Doyle 2002).

The capacitors identified by our study regulate the release of CGV across multiple environments, as demonstrated both by their ability to regulate covariance as well as by multiple epistatic interactions exhibited by them across environments. A few molecular examples of such capacitors have been identified including *HSP90* (Rutherford and Lindquist 1998), *HZT1* (Richardson *et al*. 2013) and *IRA2* (Taylor and Ehrenreich 2015). These genes have been shown to result in revelation of the hidden variants upon genetic or environmental perturbations. However, a discrepancy exists between the theoretical understanding of evolutionary capacitors and functions of these genes, *HSP90* and *HZT1,* since CGV is released both in their presence and absence. While the released CGV depends on their functional state, thus demonstrating their ability to show high epistasis, this argues against their role in evolution towards maintenance of phenotypic robustness. Such disparity in their roles questions the overlap between robustness and CGV regulation. While the capacitors identified in our study show a similar behavior, *i.e.,* both alleles have the capability to result in release of variance; they show a strong allelic bias. In majority of the cases one allele allowed the interacting locus to manifest phenotypically whereas the other allele suppressed its effect such that one allele behaved as a potentiator whereas the allele behaved as a capacitor. This along with the enrichment of such capacitors in only certain environments indicates a link between release of CGV and robustness of the phenotype.

A classic argument against an association of such loci, which regulate epistasis in standing genetic variation, with phenotypic robustness and buffering has been a lack of demonstration of their ability to accumulate new mutations. These arguments propound that CGV is independent of the state of phenotypic robustness (Paaby and Rockman 2014). While we have focused only on standing genetic variation in this study and hence cannot comment on mutational robustness, we observed a strong bias in the ability of a few environments to activate these capacitors. These capacitors have pleiotropic effects across all the environments (Cassidy *et al*. 2016) but they regulate variance only in the few environments found on the right side of the gradient and act as conventional QTL for the environments towards the left (Figure 6A). While the effect of these capacitors on novel mutations, and hence mutational robustness, cannot be demonstrated, our results confidently show that the ability of these loci to behave as capacitors is dependent on the environment and hence may be a representation of the buffered state of the environment (Fares 2015). Our results, therefore, argue in favor of evolution of these capacitors to maintain CGV (Elena and Lenski 2001).

In these specific environments, along with increased prevalence of capacitors, we identified highly interconnected networks (Figure 6B). These genetic networks were specific to each environment. The pleiotropic capacitors showed an effect in these environments but they were just one the interactors – either a part of the network or an interactor of the hub gene in the network, but never forming the core themselves (Figure 6A, B). An interesting property in these environments was the extensive cross talk that resulted in a single specific network in each environment, *i.e.,* almost all loci active in a particular environment form a part of this single network. The high number of epistatic interactions in this extreme right of the spectrum, and increased differential regulation of population variance at both single locus and two-locus levels indicates prevalence of higher order epistasis (Figure 6A, B). While we do not have the power to confidently demonstrate such high order interactions, multiple epistasic interactions between a few loci and their vQTL-vQTL mapping plots suggest their presence (see Figure 5B,C). These results support the predictions of high prevalence of such higher order interactions (Carlborg *et al*. 2006; Taylor and Ehrenreich 2014). This high interconnectivity is the reason why we observe so few additive QTL in the left side of the gradient with the majority of loci behaving as QTL+vQTL.

As described above, while the pleiotropic capacitors are also a part of these interactions, they do not form the core network. A reason for this exclusion can be that these networks could have evolved in response to specific selection pressures to reach phenotypic optimum for a particular phenotype, and the pleiotropic nature of the capacitors would implement a constraint on their ability to evolve or mold to a specific environment. These results indicate that a possible reason for the evolution of pleiotropic genes could have been the regulation of this CGV across environments. Furthermore, modularity of the interactome identified recently in disease networks (Vidal *et al*. 2011; Menche *et al*. 2015) to regulate specific diseases could be a result of evolution of these environment or phenotype specific hubs, which maintain robustness for specific phenotypes by genetic redundancy (Kafri *et al*. 2009). At a molecular level this indicates that these specific networks, which maintain robustness and contain CGV form the core regulation in each environment. A perturbation in them results in release of CGV, probably by impinging on these capacitors. Our results argue that such an interconnected system probably underlies the regulation of all complex traits but unveils itself only upon certain environmental and genetic perturbation. Different environments or phenotypes have different core networks (Chari and Dworkin 2013), which impinge on these capacitors and regulate phenotypic variation and robustness (de Visser *et al*. 2003; Chevin *et al*. 2010).

Along with high epistasis or gene-gene interactions (Mackay 2014), a major consequence of CGV is high gene-environment interactions, since these hidden variants show their effects in certain environments and not in others. While this would explain the high conditional neutrality in genetic effects across environments identified in various gene-environment interaction studies, it does not justify the observed environmental constraint on the direction of release of variance (see Figure 2B). Our results show that CGV is released in only one direction in each environment and QTL-QTL and vQTL-vQTL mapping has shown that one single network is active in each environment. Therefore, one can surmise that the hidden variation is released in only one direction by a particular network. Based on our results, we propose that a particular network, evolved to maintain optimum phenotype, can curb expression of phenotypic variation in two ways – the maximalistic approach or the minimalistic approach. The maximalistic approach would be evolution of networks that maintain high flux through the pathway(s) contributing to the phenotype such that the default state is always on. In this scenario, any genetic variant that results in a reduced flux, and hence lower phenotype, will not have much effect since there will be an excess of transcriptional or biochemical signal. As a result, such variants will get accumulated due to lack of purifying selection on them, whereas a variant that increases the flux will get fixed and become a part of the network. Perturbation of this network will, therefore, only release variants with a poorer phenotype. The alternate minimalistic approach would describe a default state with the minimum possible flux, such that the network maintains the pathway(s) in a practically shutdown state. Since the upstream signal is off or minimal, the downstream genetic variants that increase the flux will not be able to show their effects and hence get fixed. Consequently, a breakdown of this network will reveal genetic variants that increase the phenotype. Our conjecture is that modules or specialized network may have evolved this way to contain CGV in the most optimal manner. While this regulation may happen at various levels (transcription, translation or for metabolites) and differ for different phenotypes and environments, it would follow either of the two above approaches.

Along with providing molecular insights into genetic regulation of various traits, this directionality of the networks has strong implications on gene-environment interactions, especially antagonistic pleiotropy. While the core network loci are environment-specific, the generalist capacitors are common across different environments. Our results show that the abundant antagonistic pleiotropy of the mean effects of these loci is not due to opposing molecular effect of the locus on the two environments but majorly due to the opposing constraints of the environment on direction of released CGV (see Figure 3). Depending on the kind of network that impinges on the capacitor, the released CGV will either increase or decrease the phenotype. Therefore, along with regulating condition specificity of the effects of loci, regulation of CGV can result in trade-offs in the effect of loci across different environments. In our previous paper, we reported high antagonistic pleiotropy of a locus on chrXV containing the gene, *IRA2* (Yadav *et al*. 2015). *IRA2* is an inhibitor of the highly conserved Ras/PKA pathway, which we demonstrated to show high diversity in natural populations of *S. cerevisiae* adapted to diverse ecological and geographical niches. Investigation of QTL and vQTL revealed that the loci containing *IRA2* is a covariance hub and hence a capacitor (see Table S3). This is supported by a recent study, which demonstrated role of *IRA2* in regulating cryptic variation in colony morphology trait by disruption of transcriptional silencing of one or more genes that affect the trait (Taylor and Ehrenreich 2015). Comparing the mean and the variance revealed that the antagonistic effect in different environments is a consequence of differential robustness of the two alleles of *IRA2,* which may be the primary reason underlying their role in adaptation. This has profound significance on the increasing identification of antagonistic effects in various diseases causing alleles (Carter and Nguyen 2011). Our study indicates that their opposing effects are potentially a result of their differential robustness in different populations and lifestyles. This inference will be crucial in understanding their pleiotropic effects as well as their molecular mechanisms.

Our study provides a detailed understanding of the architecture of CGV affecting yeast growth across multiple environments. In doing so, we have uncovered patterns and governing principles which can be used to predict existence and behavior of CGV. This underlying architecture has the potential to elucidate certain fundamental, hitherto unclear, phenomena in quantitative biology – varying contribution of epistasis in different traits, pleiotropy, network modularity, environment and genetic background dependence of causal loci. Revisiting the already existence datasets of breeding traits and human diseases in the light of conclusions from this study can provide better insights into their genetic regulation and has the potential to solve the mystery of missing heritability (Eichler *et al*. 2010; Nelson *et al*. 2013). Identification of regulation of CGV can provide better comprehension of the molecular functioning and evolution of the modules regulating various traits and disease.

## METHODS

### Dataset

The raw growth data analysed in this study was derived from a study by Bloom *et al*. (2013), in which the experimental procedures are described in detail. The data we used was generated for 1,008 segregants derived from a cross between *S. cerevisiae* strains BY (a laboratory strain) and RM11-1a (a wine isolate, indicated as RM). These segregants were grown in 46 different conditions and phenotyped for colony size of which 34 conditions were considered in this study (see Table S1, see Files S1 and S2 for more information). A total of 11,623 markers were considered.

### QTL and vQTL mapping

The single environment QTL mapping was carried out as described previously (Bhatia *et al*. 2014). In brief, the R/qtl package (Broman *et al*. 2003; Broman and Sen 2009) was used to identify QTL separately for colony size in each environment. QTL were identified using the LOD score, which is the log10 of the ratio of the likelihood of the experimental hypothesis to the likelihood of the null hypothesis (Broman and Sen 2009). An interval mapping method (“scanone” function in R/qtl) was used to compute this LOD score using the Haley-Knott regression algorithm (Broman *et al*. 2003).

The following formula was used to calculate the F-score, which was further used to derive the LOD score. At a particular marker, let segregant i’s phenotypic value be y where *j* can take two values ( *j = 1*: BY allele and *j =* 2: RM allele).

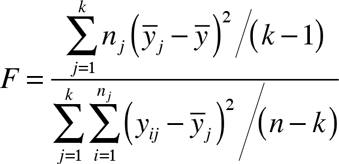

Here, *N* is the total number of segregants, *n_1_* and *n_2_* are the number of segregants having the BY and RM allele respectively ( *k =* 2) and *y_j_* is the genotypic mean of allele *j*.

Let *df* denote the degrees of freedom *(df* = 1 for a backcross and *df* = 2 for an intercross). The LOD score is accordingly derived as follows:

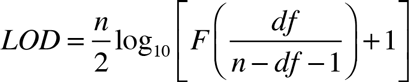

Under the null hypothesis, there is no significant difference in the means at the marker under consideration while under the alternative hypothesis, there is a presence of a QTL.

To estimate the difference in phenotypic variance between the two genotypic groups, *i.e.,* to identify vQTL in each environment, the standard Brown-Forsythe (BF) statistic (Rönnegärd and Valdar 2011; Lee *et al*. 2014) and the corresponding LOD score were calculated for each genetic marker in each environment (see Files S1 and S2). The BF test is equivalent to an F-test performed on the deviations of the phenotypic values from their respective genotypic medians (or the means). Hence, under the alternative hypothesis, the phenotypes of the two alleles reveal a difference in the variance.

At a particular marker, let *z_ij_* be the absolute deviation of segregant i’s phenotypic value *y_ij_* from its genotypic mean ÿ where *j* can take two values ( *j = 1*: BY allele and *j* = 2: RM allele).

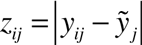

Then BF statistic for that marker can be computed as follows:

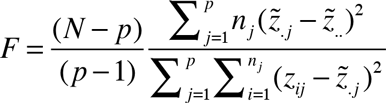

Here, *N* is the total number of segregants, *n_1_* and *n_2_* are the number of segregants having the BY and RM allele respectively ( *p* = 2). In order to estimate the effects of vQTL in the same order as in QTL, LOD scores were computed as described previously (Broman and Sen 2009).

To establish the statistical significance of the putative QTL and vQTL, *P* values were computed using a genome-wide permutation test of 1,000 permutations, where the null distribution consisted of the highest genome-wide LOD score obtained from each permutation (Broman *et al*. 2003). A LOD score cut off of greater than 3.0 and a *P* value cut off of less than 0.01 was considered.

### Comparing QTL and vQTL

To estimate pleiotropy, we divided the genome into 20kb non-overlapping bins (Table S3). The bins containing two or more QTL or vQTL significant (P < 0.01) in different environments were considered as pleiotropic bins. The first markers of each of these pleiotropic bins, used as representative of the bins, were collated to represent the set of pleiotropic markers (Table S3).

### Calculating correlation between mean and variance

To calculate environment-specific correlation (for Figure 2C), difference in the mean (mean BY – mean RM) and difference in the variance (var BY – var RM) was calculated for all significant loci within each environment. Pearson’s correlation between these two parameters was calculated for each environment.

### Covariance across environmental pairs

To assess the differential covariance of a locus across multiple pairs of environments, we considered the collated set of pleiotropic markers for our study (Table S3). To quantify the differential covariance across a pair of environments, a Deming regression was calculated between the phenotype values of the chosen pair of environments for each allele, using R package ‘mcr’. Deming regression, which minimizes errors in multiple dimensions simultaneously, served as a suitable measurement error model for assessing buffering across two or more environments. Unlike simple least squares regression, Deming regression accounts for deviations in observations on both the *x*- and the *y*- axis. It seeks to find the line of best fit

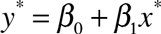

such that the below weighted sum of squared residuals is minimized,

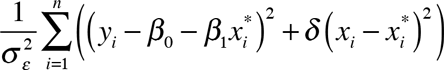

where, the estimate of *δ* is the ratio of the two variances.

Since both the phenotypic values (*x* and *y*) are normally distributed across their respective axes, Deming regression, serves as a suitable measurement-error model for assessing buffering across two environments. Two Deming regression models were fitted for each environment pair corresponding to the two alleles.

A Student’s t-test was performed between the deviations of the phenotypic values from the Deming fit of the BY and RM alleles (P < 0.05). This sign of the t-test value represented the allele that had higher mean of deviations, *i.e.,* lower covariance. If t-test value was positive, then the mean deviation of BY allele was greater than that of the RM allele and vice versa. All the significant markers and their corresponding environmental pairs along with mean deviations of the alleles and independent mean and variance of both the alleles and both the environments are listed in Table S3. A locus with significantly different allelic covariance across more than 15 environmental pairs was considered pleiotropic. For each of these loci, total number of positive and negative t-test statistic were compiled. A Fisher’s Exact test of random distribution of positive and negative t-test statistic was done each locus, normalized for the number of environmental pairs. A *P* value cut-off less than 0.05 was considered significant.

### QTL-QTL and vQTL-vQTL mapping

A QTL-QTL interaction occurs when an effect of a QTL at a single locus depends on another locus. We used a QTL-QTL mapping technique described previously (Bhatia *et al*. 2014). In brief, we used a custom Python script, from Bhatia *et al*. (2014) to compute LOD scores for pairwise comparisons among a set of selected markers. These markers were selected using a QTL *P* value cut-off of 0.01 in each environment. For the QTL-QTL mapping, *P* values were computed using a permutation test (10,000 permutations), where the null distribution consisted of the highest LOD score obtained among all pairwise comparison for each permutation of the phenotype. The following hypotheses were compared:

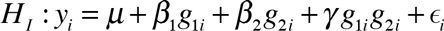

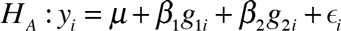

Here, *g*_1_*i* and *g*_2_*i* are the binary variables that specify the genotype at the two loci, *μ, β*_1_, *β_2_* and *γ* are inferred from the data using maximum likelihood. The parameters *β*_χ_ and *β*_2_ quantify the individual effect of each QTL, and *γ* quantifies the effect of the QTL-QTL interaction. See Files S1 and S2 for details of calculation of the interaction LOD ( *LOD_i_*) score.

Apart from the QTL-QTL mapping described previously (Bhatia *et al*. 2014), we mapped variance-controlling interactions by mapping vQTL-vQTL interactions, which occurs when the phenotypic variance at one locus depends on the genotype at another locus. The hypothesis testing for vQTL-vQTL interactions was done in the same way as QTL-QTL interactions except that instead of scaled values of colony size as the phenotypic values, deviations from the mean were used as the phenotype for each allelic combination (see method for vQTL mapping).

To increase power to identify QTL-QTL and vQTL-vQTL interactions, for each environment, genetic loci significant in either QTL or vQTL or QTL+vQTL mappings were collated for each environment (Table S1). To compile QTL-QTL and vQTL-vQTL interaction results, loci within a 50kb linkage interval were considered the same. The *P* values were computed using a permutation test of 10,000 permutations with the phenotype data shuffled relative to the genotype data.

## ACKNOWLEDGEMENTS

This research was supported by Tata Institute of Fundamental Research intramural funds (HS). The funders had no role in study design, data collection and analysis, decision to publish, or preparation of the manuscript.

## AUTHOR CONTRIBUTIONS

Conceived and designed analysis: AY HS; Analysed data: KD AY HS; Wrote the paper: AY

## SUPPORTING INFORMATION LEGEND

**File S1**: Scripts and datasets for QTL, vQTL, Deming regression and QTL-QTL, vQTL-vQTL interaction mapping.

**File S2**: Supplementary methods.

**Table S1**: QTL and vQTL mapping

QTL test and vQTL test refers to the F-test and BF-test statistics respectively, and QTL *P* value and vQTL *P* value refers to the *P* values of the corresponding tests. Test cutoff of >3 and *P* value cutoff of <0.005 was used to identify significant markers. 4-HBA refers to 4-Hydroxybenzaldehyde; 4-NQO is 4-Nitroquinoline; 5-FC is 5-Fluorocytosine; 5-FU is 5 Fluorouracil; 6-AU is 6-Azauracil.

**Table S2**: Mean and variance of significant QTL and QTL+vQTL

**Table S3**: t-test of Deming regression to compare trait covariance

t-test refers to t-test statistic of Student’s t-test performed on Deming regression values of the two alleles of a maker across environment 1 and 2. meanRM and meanBY refers to mean of the regression values of RM and BY alleles, respectively, across environment 1 and 2. meanRM_env1 and varRM_env1refers to mean and variance of the RM allele in environment 1. Same nomenclature applies to other alleles and environments.

**Table S4**: QTL-QTL and vQTL-vQTL mapping

Permutation *P* value cutoff of < 0.1 was considered. Sheet 1 contains results of QTL-QTL mapping and Sheet 2 contains results of vQTL-vQTL mapping. Sheet 3 shows the overlap of the Sheet 1 and Sheet 2. In Sheet 3, markers with a 50kb interval were considered the same to estimate the overlap between the two kinds of mapping. A significant QTL-QTL or a vQTL-vQTL interaction is indicated by 1.

**Table S5**: Classifying interacting loci in specific gene-gene interactions, capacitors and environment-specific genetic networks.

